# Keloid transcriptomics reveal heterogeneity in fibroblast subtype enrichment, gene expression, and immune cell responses

**DOI:** 10.64898/2026.09.02.747090

**Authors:** Jonathan J. Panzer, Mingming Pan, Mrudula Nair, Ian M. Loveless, Xiangguo Dai, Indra Adrianto, Ling Huang, Dhananjay Chitale, Ralph Francescone, Debora B. Vendramini-Costa, Cristina de Guzman Strong, Albert M. Levin, Lamont R. Jones

**Affiliations:** Department of Public Health Sciences, Henry Ford Health, Detroit, Michigan, USA; Department of Otolaryngology, Henry Ford Medical Group, Detroit, Michigan, USA; Henry Ford Health + Michigan State University Health Sciences, Detroit, Michigan, USA; Department of Medicine, Michigan State University, Lansing, Michigan, USA; Henry Ford Pancreatic Cancer Center, Surgery, Detroit, Michigan, USA; Department of Pharmacology and Toxicology, Michigan State University, Lansing, Michigan, USA; Department of Pathology, HFH, Detroit, Michigan, USA; Department of Pathology, Michigan State University, Lansing, Michigan, USA; Department of Physiology, Michigan State University, Lansing, Michigan, USA; Department of Dermatology, HFH, Detroit, Michigan, USA; Department of Epidemiology and Biostatistics, Michigan State University, Lansing, Michigan, USA; Department of Otolaryngology, Michigan State University, Lansing, Michigan, USA

**Keywords:** Fibroproliferative, abnormal wound healing, fibrosis, cellular programming, African American

## Abstract

Keloid disease (KD) is a fibroproliferative skin disorder resulting from abnormal scar formation that causes pain, itching, and decreased quality of life. While multiple KD transcriptomic studies exist, the influence of cell type composition on bulk tissue gene expression is unknown. We characterized fibroblast subtype and immune cell enrichment using bulk RNA-Seq of head and neck keloid and matched adjacent normal skin tissue (MANST) from 14 patients (10 African American and 4 European American). Cell type enrichment was calculated by single sample gene set enrichment analysis. Linear mixed-effects models were employed for 1) differential cell type enrichment across tissue, 2) tissue type-specific associations between fibroblast subtypes and immune cells, and 3) differentially expressed genes (DEGs) across tissue. Validation was conducted in an independent cohort of 8 African Americans. Three fibroblast subtypes and 14 immune cell types were differentially enriched across tissue type. Further, 17 tissue type-specific fibroblast subtype-immune cell enrichment associations were identified, with 14 exhibiting decreased association in keloid tissue relative to MANST. After adjustment for cell type enrichment, MIR31HG and NR4A2 were significant DEGs with the largest positive and negative fold-changes, respectively. By considering cell type enrichment, underlying keloid tissue-specific cell type and gene expression associations were revealed.

## INTRODUCTION

Keloid disease (KD) is a fibroproliferative disorder involving abnormal wound healing that leads to excessive collagen deposition, persistent inflammation, and overgrowth of scar tissue beyond the original wound (Alonso et al. 2008; Ashcroft et al. 2013; Deng et al. 2021; Fang et al. 2025). KD patients suffer from pain, disfigurement, and pruritus that negatively impacts their quality of life (Bijlard et al. 2017). Unlike normal scars, keloids do not resolve on their own and often recur after treatment (Gold et al. 2020). While keloids disproportionately impact African Americans, Asians, and Hispanics (Swenson et al. 2024), African Americans have been underrepresented in molecular studies of KD. Yet, inclusion of African Americans and other minority populations impacted by KD is essential to increase our understanding of the heterogeneity of keloid tissue dysregulation. Ultimately, enabling progress towards the precise selection of existing treatments and/or the development of novel KD-specific therapeutics that will improve treatment outcomes in a broader range of individuals.

Fibroblasts that over-produce extracellular matrix (ECM) components (e.g., collagen) are central to keloid development and progression (Ashcroft et al. 2013; Liu et al. 2022). However, fibroblast heterogeneity and its impact on keloid biology is only starting to be explored. For example, a recent single cell RNA sequencing (scRNA-Seq) study of non-diseased, sun-protected fair skin identified five fibroblast subtypes (Solé-Boldo et al. 2020), and initial KD studies indicate that the abundance of the mesenchymal fibroblast subtype is a significant differentiator between keloid and normal skin tissue (Mao et al. 2024; Shim et al. 2022; Xia et al. 2023; Xie et al. 2025; Yeo et al. 2024; Zhao et al. 2025). However, it is not known whether this is true in other populations such as African Americans, who have the highest KD incidence and more aggressive disease (Margiotta et al. 2022; Olopoenia et al. 2024; Robles and Berg 2007; Sharma et al. 2025; Velez Edwards et al. 2014; Young et al. 2014).

While fibroblasts are central to keloid scars, they are not the only cell type participating in keloid pathology. Many immune cell populations (e.g., mast cells, M2 macrophages, and T cells) also contribute to the abnormal maintenance of the wound healing state in keloid tissue (Liu et al. 2026). These immune populations interact with fibroblasts to reinforce fibroblast activation, sustain chronic inflammation and fibrosis, and disrupt normal resolution (Li et al. 2017; Nakajima et al. 2022; Robles and Berg 2007; Yeo et al. 2024). Despite the contribution of immune cell populations to keloid scars, few studies have investigated how fibroblast subtype composition may differentially influence the immune cell composition in keloid relative to normal skin. Further, transcriptomic studies comparing keloid tissue to normal skin have typically not accounted for fibroblast and immune cell composition, resulting in differential gene findings that may be confounded by the differential enrichment of these cell types.

To address these KD knowledge gaps, we performed bulk RNA sequencing (RNA-Seq) on head and neck keloid tissue and matched adjacent normal skin tissue (MANST) from a majority African American patient cohort. First, we characterized how fibroblast subtype and immune cell composition differ across keloid tissue and MANST. Next, we explored how the enrichment between fibroblast subtype-immune cell pairs differs in keloid tissue and MANST. Finally, we utilized the knowledge gained from the preceding cell type composition analyses in an assessment of differential gene expression between keloid tissue and MANST. To ensure the reliability of our findings from these three analyses, we validated key observations with data from a previously published RNA-Seq study that included African American paired keloid tissue and MANST (Zhu et al. 2023).

## RESULTS

### Keloid cohort description

Patient-paired keloid tissue and matched adjacent normal skin tissue (MANST) were obtained from 14 patients with primary, previously untreated keloids in the head and neck (H&N) region. These patients were all undergoing keloid removal surgery within the Department of Otolaryngology at Henry Ford Health (HFH; Detroit, Michigan). Characteristics of the cohort are summarized in **Table 1**. Patient age at surgery varied from 15 to 86 years; a majority of the cohort was African American (n=10, 71.4%); and eight (57%) of the 14 participants were female (**Table 1**). Additionally, 13 of the 14 H&N keloids (93%) were from the ear.

**Table 1.** Summary statistics for the HFH keloid patient cohort (n = 14).

| **Characteristic** | **Levels** | **Overall** |
| --- | --- | --- |
| **Age (mean +/- SD)** |  | 38.4 (23.3) |
| **Race (n (%))** | **African American** | 10 (71.4) |
|  | **European American** | 4 (28.6) |
| **Sex (n (%))** | **Female** | 8 (57.1) |
|  | **Male** | 6 (42.9) |
| **Keloid Location (n (%))** | **Ear** | 13 (92.9) |
|  | **Neck** | 1 (7.1) |

### Differential enrichment of fibroblast subtypes and immune cells between keloid tissue vs. MANST

To investigate the potential differential contribution of fibroblast subtypes and immune cell types to keloid tissue and MANST, ssGSEA-derived fibroblast subtype (n=5) and immune cell (n=22) enrichment scores were calculated for each specimen. In addition to providing insight into differences in cellular heterogeneity between the two tissue types, this assessment was also critical as differences in cell type and subtype abundances between the tissue types could confound subsequent differential gene expression analyses. Three out of the five fibroblast subtypes were significantly enriched in keloid tissue relative to MANST: secretory-reticular (p=9.55*10^-6^), mesenchymal (p=3.08*10^-5^), and pro-inflammatory B (p=0.003) (**Figure 1, Figure S1**). Mesenchymal (p=0.006) and pro-inflammatory B (p=0.003) fibroblast subtypes were also significantly enriched in keloid tissue relative to MANST in the Zhu et al. dataset (**Table S1**).

**Figure 1.**
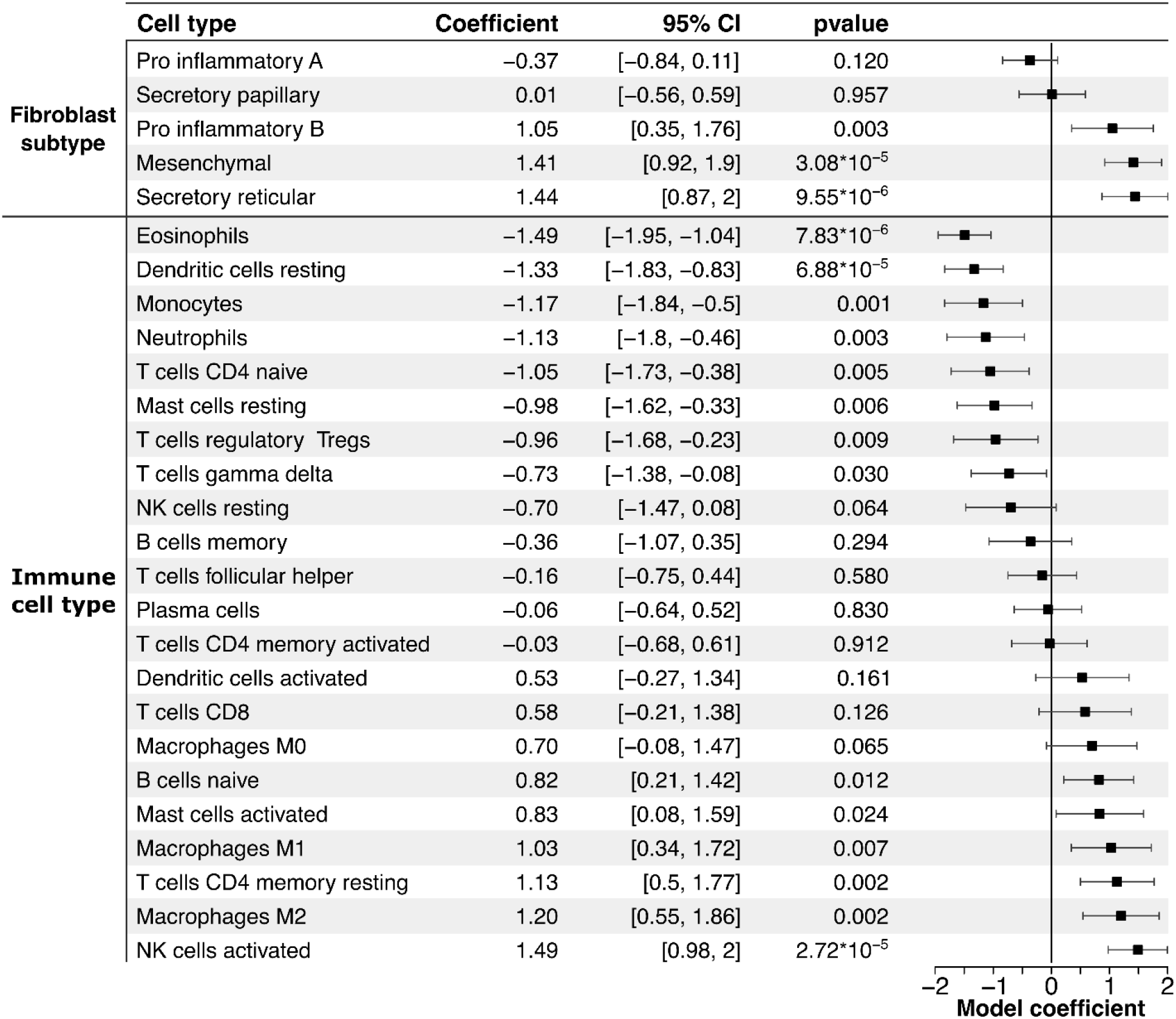
Forest plot of fibroblast subtype and immune cell enrichment score associations with keloid tissue versus MANST. Model coefficient and 95% confidence intervals are displayed from linear mixed models testing for enrichment scores differences between keloid tissue and MANST (i.e., positive model coefficients are indicative of enrichment of a particular cell type in keloid tissue relative to MANST, while negative model coefficients reflect the opposite relationship). As the enrichment scores for each cell type have been normalized to have a mean value of zero and a standard deviation (and variance) of one, the model coefficients are interpretable in standard deviation unit differences between the two tissue types (e.g., the mean mesenchymal fibroblast score is 1.41 standard deviation units higher in keloid tissue vs. MANST). Within each group of cell types, results are displayed in ascending model coefficient order.

Across the 22 immune cell types, six were significantly enriched and eight were significantly diminished in keloid tissue relative to MANST (**Figure 1, Figure S1**). More specifically, both differentiated macrophage subtypes, M1 (p=0.007) and M2 (p=0.002), were increased in keloid tissue. In addition, enrichment of activated mast cells (p=0.024) and NK cells (p=2.72*10^-5^) was also increased in keloid tissue relative to MANST, whereas the resting types of these cell populations were diminished (resting mast cell, p=0.006; resting NK cell, p=0.064). Lastly, CD4 memory resting T cells (p=0.002) and naïve B cells (p=0.012) were the only other immune cell types significantly enriched in keloid tissue. In contrast, gamma delta T cells (p=0.030), CD4 naïve T cells (p=0.005), neutrophils (p=0.003), resting dendritic cells (p=6.88*10⁻⁵), and eosinophils (p=7.83*10⁻⁶) were significantly diminished in keloid tissue relative to MANST. Similarly, the Zhu et al. dataset revealed significant keloid tissue enrichment for both differentiated macrophage subtypes M1 (p=7.29*10^-4^) and M2 (p=0.001), as well as activated mast cells (p=9.02*10^-4^) and activated NK cells (p=0.002) (**Table S1**).

### Tissue type-specific associations between fibroblast subtype and immune cell enrichment

To gain further insight into how fibroblast subtype composition may be associated with immune cell composition in keloid tissue relative to MANST, associations between enrichment scores for fibroblast subtype and immune cell pairs were evaluated across tissue type. Those pairs with significant differences in associations between keloid tissue and MANST (p_interaction_ <0.1) were highlighted in **Figure 2**, with all findings summarized in **Table S2**. Positive associations reflect concordant increase (or decrease) in immune cell enrichment associated with increase (or decrease) in fibroblast subtype enrichment, while negative associations reflect discordant (i.e., oppositely directed) association between the fibroblast subtype-immune cell pair. A total of 17 tissue type-specific pair associations were identified, and pairs including secretory-reticular (n=8 pairs) and mesenchymal fibroblasts (n=6 pairs) accounted for the majority. The tissue type-specific associations (keloid association = β_keloid_; and MANST association = β_MANST_) could be categorized into two distinct patterns: 1) β_keloid_ > β_MANST_ (n=3 pairs) and 2) β_keloid_ < β_MANST_ (n=14 pairs). For pattern #1, the most significant interaction was observed between secretory-reticular fibroblasts and activated natural killer (NK) cells (β_keloid_ = 0.92, p-value=0.008; β_MANST_ = −0.05, p-value=0.723; p_interaction_= 0.004). For pattern #2, the most significant interaction was observed between secretory-reticular fibroblast and eosinophils (β_keloid_ = −0.89, p-value=0.017; β_MANST_ = 0.06, p-value=0.510; p_interaction_= 0.006).

**Figure 2.**
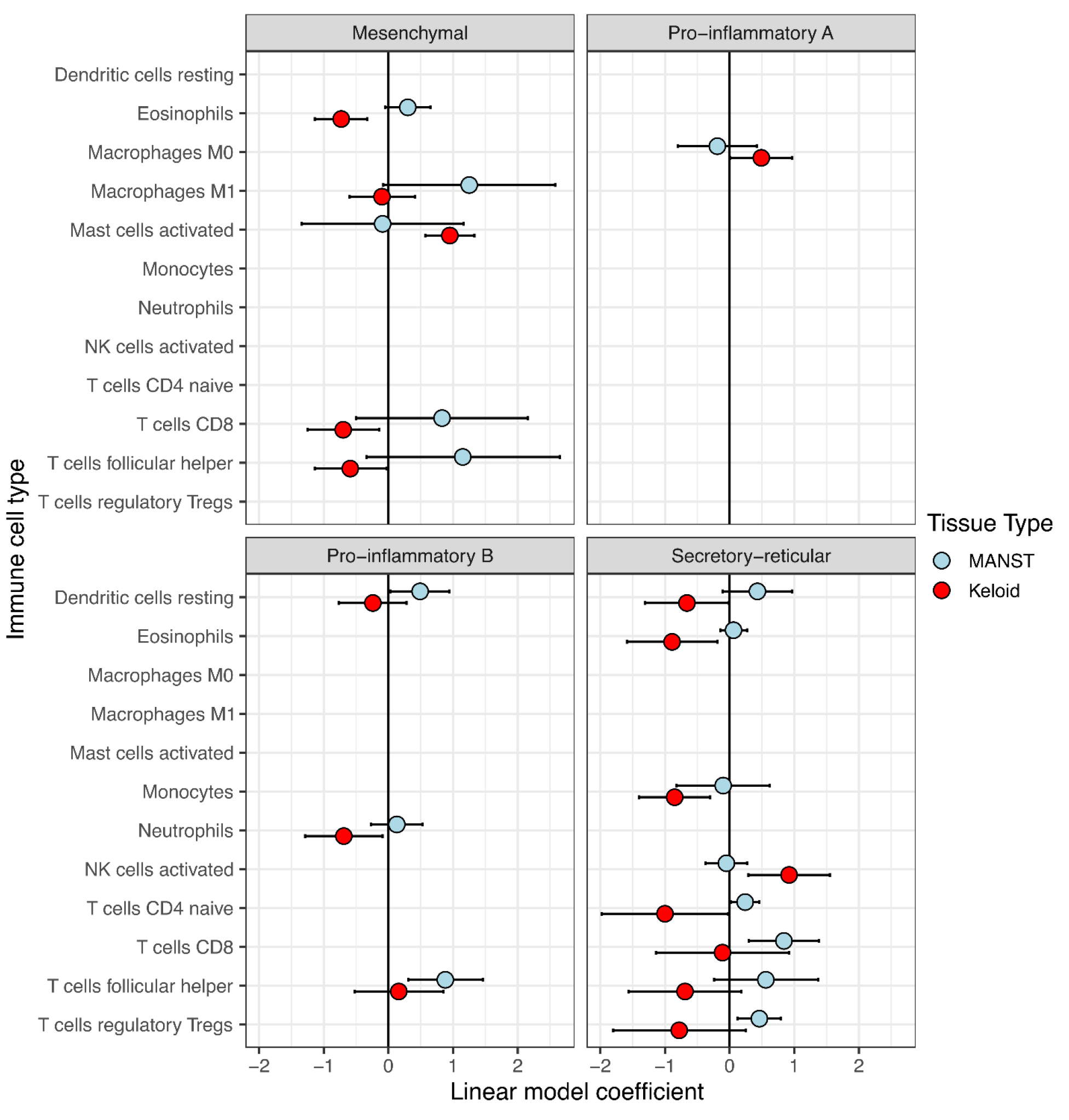
Fibroblast subtype-immune cell associations differing by tissue type. Linear model coefficients (boxes – keloid, circles - MANST) and 95% Wald confidence intervals (bars) plotted for tissue type stratified models. Only fibroblast subtype-immune cell associations that differed between keloid tissue and MANST (interaction term p-value < 0.1) were included.

The Zhu et al. dataset also revealed a significant interaction between secretory-reticular fibroblasts and activated natural killer (NK) cells (β_keloid_ = 0.33, p-value=0.23; β_MANST_ = −0.33, p- value=0. 23; p_interaction_= 0.098) (**Table S3**). While the remaining 16 interactions were not significant in the Zhu et al. dataset, 12 of the 17 pairs matched the tissue type coefficient patterns originally identified (**Table S3**).

### Keloid tissue vs. MANST differentially expressed gene (DEGs) and biological pathway analyses

Next, analyses were performed to identify DEGs between keloid tissue and MANST. After quality control, 15,753 genes were available for analysis. Given the tissue-specific fibroblast subtype and immune cell compositions described above, DEG tests were adjusted for the enrichment scores of these cell types to mitigate potential confounding. Specifically, principal components analysis (PCA) was performed on the combined fibroblast subtype and immune cell enrichment scores (i.e., 27 cell type enrichment scores: 5 fibroblast subtypes and 22 immune cells) to reduce the dimensionality of the data. The top three principal components from this PCA (accounting for 62.3% of the total variation of the enrichment scores) were adjusted for in the analyses (**Figure S2**). The transcriptome-wide results are presented in **Table S4**. Overall, 1,145 genes were nominally significant (p-value < 0.05), but only 15 genes were differentially expressed after FDR correction (FDR-adjusted p < 0.05) (**Table 2**). Notably, among the 15 FDR-associated genes, the gene with the largest positive fold-change between keloid tissue and MANST was MIR31HG (MIR31 Host Gene; log2 fold-change=3.41, FDR-adjusted p=3.08*10⁻^5^), while the largest negative fold-change was observed for NR4A2 (Nuclear Receptor Subfamily 4 Group A Member 2; log2 fold-change=-1.59, FDR-adjusted p=3.08*10⁻^5^). Four of the 15 FDR-associated genes were also nominally significant with concordant directions of effect in the Zhu et al. dataset including MIR31HG (p=0.013), GJC1 (p=0.004), CHN1 (p=0.029), and ETV1 (p=0.001) (**Table S5**). Overall, 12 of the 15 DEGs maintained directionality across both datasets (**Table S5**).

**Table 2.**
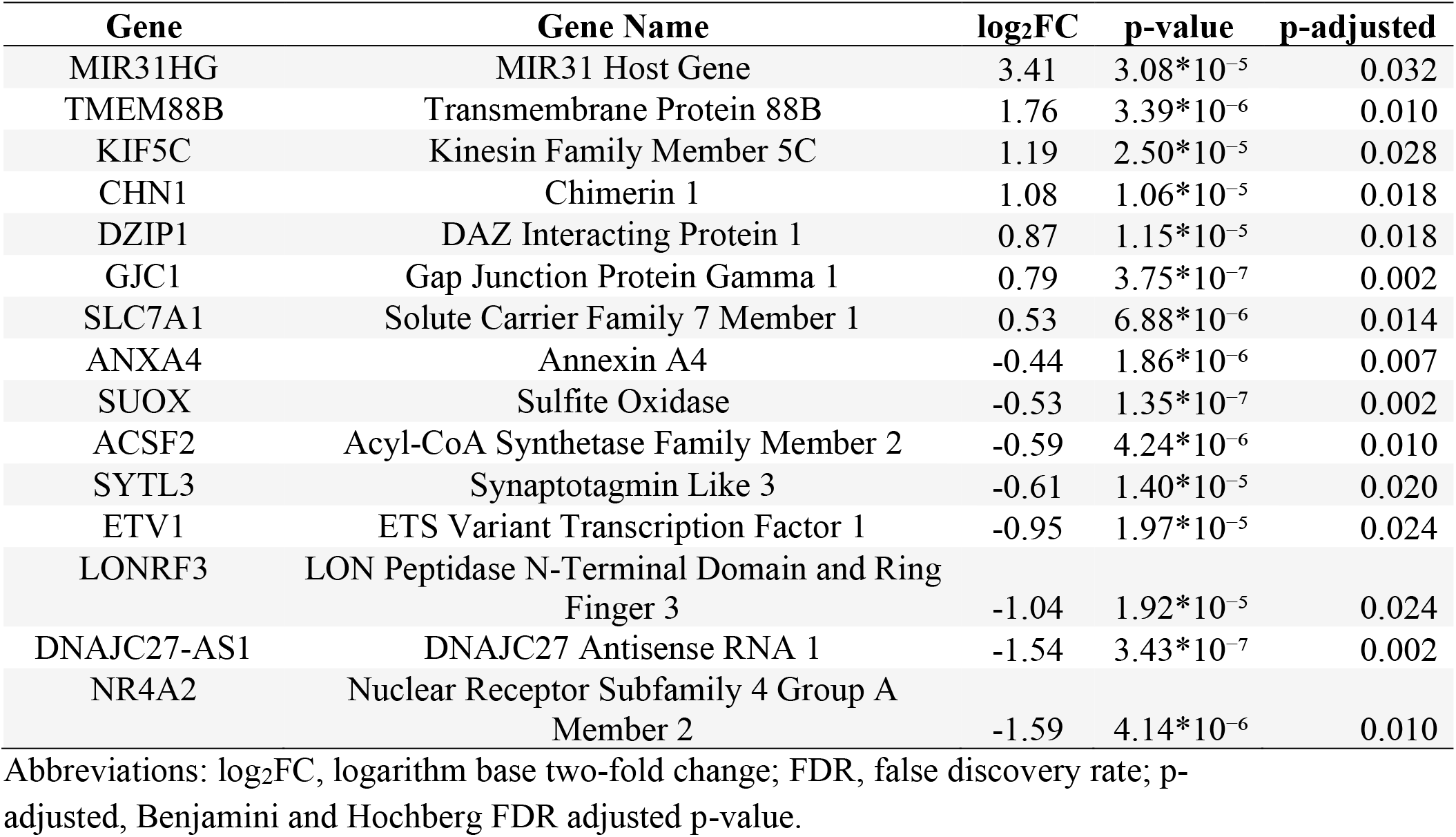
FDR-significant DEGs across tissue type. DEGs were sorted by log_2_FC after filtering for significant DEGs (FDR-adjusted p< 0.05), with MANST expression as reference.

| **Gene** | **Gene Name** | **log<sub>2</sub>FC** | **p-value** | **p-adjusted** |
| --- | --- | --- | --- | --- |
| MIR31HG | MIR31 Host Gene | 3.41 | 3.08*10<sup>-5</sup> | 0.032 |
| TMEM88B | Transmembrane Protein 88B | 1.76 | 3.39*10<sup>-6</sup> | 0.010 |
| KIF5C | Kinesin Family Member 5C | 1.19 | 2.50*10<sup>-5</sup> | 0.028 |
| CHN1 | Chimerin 1 | 1.08 | 1.06*10<sup>-5</sup> | 0.018 |
| DZIP1 | DAZ Interacting Protein 1 | 0.87 | 1.15*10<sup>-5</sup> | 0.018 |
| GJC1 | Gap Junction Protein Gamma 1 | 0.79 | 3.75*10<sup>-7</sup> | 0.002 |
| SLC7A1 | Solute Carrier Family 7 Member 1 | 0.53 | 6.88*10<sup>-6</sup> | 0.014 |
| ANXA4 | Annexin A4 | -0.44 | 1.86*10<sup>-6</sup> | 0.007 |
| SUOX | Sulfite Oxidase | -0.53 | 1.35*10<sup>-7</sup> | 0.002 |
| ACSF2 | Acyl-CoA Synthetase Family Member 2 | -0.59 | 4.24*10<sup>-6</sup> | 0.010 |
| SYTL3 | Synaptotagmin Like 3 | -0.61 | 1.40*10<sup>-5</sup> | 0.020 |
| ETV1 | ETS Variant Transcription Factor 1 | -0.95 | 1.97*10<sup>-5</sup> | 0.024 |
| LONRF3 | LON Peptidase N-Terminal Domain and Ring<br>Finger 3 | -1.04 | 1.92*10<sup>-5</sup> | 0.024 |
| DNAJC27-AS1 | DNAJC27 Antisense RNA 1 | -1.54 | 3.43*10<sup>-7</sup> | 0.002 |
| NR4A2 | Nuclear Receptor Subfamily 4 Group A<br>Member 2 | -1.59 | 4.14*10<sup>-6</sup> | 0.010 |

Abbreviations: log<sub>2</sub>FC, logarithm base two-fold change; FDR, false discovery rate; p-adjusted, Benjamini and Hochberg FDR adjusted p-value.

IPA-based canonical biological pathway analysis of the 1,145 nominally significant (p<0.05) DEGs uncovered gene enrichment for 62 pathways (p<0.001), and the complete pathway analysis results are presented in **Table S6**. We further evaluated these enriched pathways based on the predicted regulation of the pathways using the IPA pathway specific activation Z-scores. At an absolute activation Z-score greater than two (i.e., > 2 indicating upregulation or < −2 indicating down regulation), 18 pathways consistent with keloid fibrotic pathology were revealed (**Figure 3A**), all of which were predicted to be upregulated in keloid tissue vs. MANST. These pathways included “Pulmonary Fibrosis Idiopathic Signaling Pathway”, “Hepatic Fibrosis Signaling Pathway”, “Collagen Biosynthesis and Modifying Enzymes”, and “Collagen degradation”. Additionally, activation of several pathways suggested inflammatory activation including the “S100 Family Signaling” and “Eicosanoid Signaling” pathways. Further, predicted activation of the “Molecular Mechanisms of Cancer” pathway emphasized commonalities between tumor and keloid growth (**Figure 3A**). An IPA graphical synthesis of relationships between canonical pathways, cellular functions, and DEGs, highlighted the predicted activation of TNF (Tumor Necrosis Factor) along with NRG1 (Neuregulin 1) and multiple interferons (IFNL1, IFNA2, and IFNG), leading to tumor-related cellular functions, cell growth and differentiation, and inflammation (**Figure 3B**).

**Figure 3.**
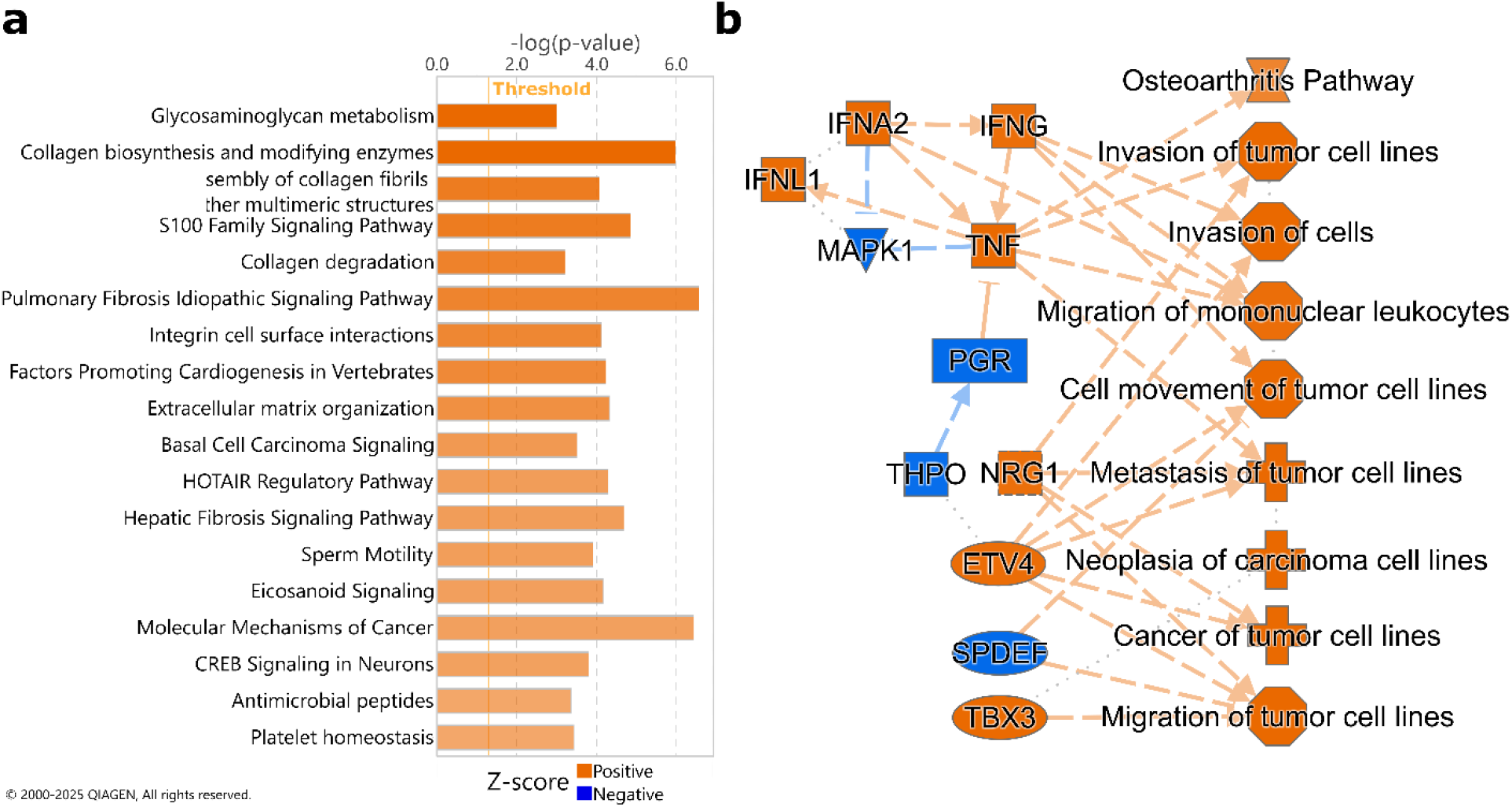
IPA canonical pathway enrichment and graphical summary based on tissue type-associated DEGs. A) Canonical pathways sorted by activation z-score after filtering for significance (-log_10_(p) > 3) and absolute z-score > 2. B) Graphical summary of relationships between and predicted activation state of canonical pathways, cellular functions, and DEGs.

## DISCUSSION

Fibroblasts are of critical importance to keloid biology and are well known to be heterogeneous in their gene expression, leading to versatile fibroblast populations with mixed functionalities (Macarak et al. 2021; Steele et al. 2025). To untangle this fibroblast heterogeneity in keloid tissue, multiple studies have attempted to characterize fibroblast subtypes through scRNA-Seq, with the number of identified fibroblast subtypes ranging from 3 to 7 (Cheng et al. 2024; Deng et al. 2021; Gong et al. 2022; Liu et al. 2022; Mao et al. 2024; Shim et al. 2022; Xie et al. 2025; Zhao et al. 2025). To explore the impact of fibroblast heterogeneity in the current study, we adopted fibroblast subtyping nomenclature from a study of clinically healthy skin that categorized fibroblasts into 5 subtypes (Solé-Boldo et al. 2020). Through the resulting enrichment analyses, we demonstrated that keloid tissue is enriched for mesenchymal, secretory-reticular, and pro-inflammatory B fibroblast subtypes. Indeed, mesenchymal fibroblasts, originally described as fibroblasts with a greater potential to differentiate into other mesenchymal cell types, are widely recognized to be more enriched in keloid tissue when compared to normal scar (Deng et al. 2021), matched non-lesional (Gong et al. 2022; Liu et al. 2022), and normal, non-patient matched skin tissue (Shim et al. 2022; Zhang et al. 2025; Zhao et al. 2025). Intriguingly, all three keloid-enriched fibroblast subtypes were originally identified as localized to the reticular dermis (Solé-Boldo et al. 2020), implying that the aberrant wound healing process leading to keloid development may be especially apparent in the deeper dermis.

Additionally, our findings highlighted immune cell abundance associations with keloid tissue including activated mast cells, activated NK cells, and differentiated macrophages (both M1 and M2). Mast cells are known to be increased in keloid tissue (Tang et al. 2023; Xiao et al. 2024; Zhu et al. 2023), and activated mast cells promote fibrosis and stimulate fibroblast activation (Ammendola et al. 2013; Liu et al. 2026). Activated NK cells were similarly enriched in keloid tissue, yet their role in keloid pathogenesis is unclear given their typical connection with antiproliferative cytokine production and cytotoxicity. However, a recent study demonstrated that keloid-associated NK cells were of an exhausted phenotype, thus nullifying their typical antiproliferative role (Zhao et al. 2026). Differentiated macrophages are also known to be increased in keloid tissue as opposed to healthy tissue. Specifically, CD163+ M2 macrophages exhibit markedly increased infiltration within keloid tissue in comparison to CD68+ M1 macrophages (Li et al. 2017), skewing the macrophage response towards fibrosis (Strizova et al. 2023). Notably, eosinophils, responsible for combating infections and some allergic responses (Wechsler et al. 2021), were the most diminished immune cell type in keloid tissue.

Given the well-established role of fibroblasts in coordinating immune signaling during wound healing, we also explored the potential for differential association between fibroblast subtypes and immune cells between keloid tissue and MANST. Of the two patterns: β_keloid_ > β_MANST_ and β_keloid_ < β_MANST_, the latter accounted for nearly all, indicating mostly decreased associations between fibroblast subtypes and immune cells in keloid tissue. The most significant interaction matching the latter pattern (β_keloid_ < β_MANST_) was between secretory-reticular fibroblasts and eosinophils. While eosinophil-fibroblast interactions are known to induce fibrosis (Gomes et al. 2005), the observation of a weaker association in keloid tissue may reflect the chronic fibrotic state of keloid tissue rather than eosinophilic stimulation of fibroblasts that characterizes the earlier stages of wound healing (Coden and Berdnikovs 2020; Todd et al. 1991). In contrast, one of the strongest positive keloid tissue-specific associations (i.e. not evident in MANST) was activated mast cells with mesenchymal fibroblasts. Mast cells have been observed in close proximity to myofibroblasts with direct cell-cell contacts in keloid tissue (Beer et al. 1998; Macarak et al. 2021), and both mast cells and mesenchymal-like fibroblasts are implicated in keloid disease progression. Together with previous findings, these results suggest either that mast cells preferentially associate with mesenchymal and secretory reticular fibroblast subtypes or that mast cell–driven stimulation directs fibroblasts toward mesenchymal or secretory reticular states. Additionally, activated NK cells were not only found to be associated with keloid tissue but were also found to have a keloid tissue-specific association with secretory reticular fibroblasts, implying that NK cell dysregulation may lead to uncontrolled proliferation of the secretory reticular fibroblast subtype in KD. Further, all tissue specific fibroblast-T cell associations exhibited a decoupling in keloid tissue compared to MANST, with the most shifts occurring for secretory-reticular and mesenchymal fibroblast subtypes. One of these, CD8+ T cells, which can inhibit proliferation of keloid-associated fibroblasts (Shan et al. 2022; Zhang et al. 2023b), exhibited decreased association with both mesenchymal and secretory-reticular fibroblast subtypes in keloid tissue.

Given the complex cell type composition differences between keloid tissue and MANST, a differential gene expression analysis across tissue type would have been severely confounded had we not directly accounted for fibroblast and immune cell composition, which is a known concern more broadly (Jin et al. 2021). Specifically, cell type composition is known to influence observed differences in gene expression across tissues or conditions, often obscuring the true underlying transcriptional changes that shape the tissue microenvironment and drive disease processes such as keloid formation (Repsilber et al. 2010; Shen-Orr et al. 2010). This is particularly important when the proportions shift significantly across tissue type for cell types contributing to the observed differential phenotype (Deng et al. 2021; Limandjaja et al. 2020; Ogawa 2017). Therefore, it is essential to adjust for these compositional differences by incorporating cell type proportions or enrichment scores as covariates in statistical models for gene expression. While three studies on bulk keloid tissue performed either bulk deconvolution (Zhu et al. 2023) or ssGSEA (Tang et al. 2023; Xiao et al. 2024), no other published bulk RNA-Seq study of DEGs across keloid and normal skin tissue has applied adjustment for cell type composition.

After adjustment for cell type composition, the MIR31 host gene (MIR31HG) had the largest over-expression in keloid tissue relative to MANST. MIR31HG encodes a long non-coding RNA (lncRNA) implicated in multiple biological processes that are dysregulated in keloids, including the cell cycle, cell proliferation, epithelial to mesenchymal transition, apoptosis, inflammation, and senescence (Ko and Yang 2025; Kolenda et al. 2023). Additionally, MIR31HG also encodes for a microRNA, miR31, which can be generated from an intronic region of the MIR31HG primary transcript through canonical microRNA processing. In a previous study, we observed differential miR31 expression in keloid tissue relative to MANST, and miR31 belongs to a diagnostic biomarker profile composed of 17 microRNA that discriminates between the two tissue types (Jones et al. 2024). In a subsequent study, we also found that miR31 expression in keloid tissue is associated with recurrence risk following surgical removal of a primary, previously untreated keloid (Levin et al. 2024). Taken together, these results suggest that the lncRNA gene MIR31HG may play an important role in both keloid development and aggressiveness.

The Nuclear Receptor Subfamily 4 Group A Member 2 (NR4A2) gene was the most downregulated DEG in keloid tissue relative to MANST. NR4A2 is an orphan nuclear receptor with no known endogenous ligand that acts as an inducible transcription factor by binding promoters of target genes with NBRE motifs (as a monomer or homodimer), NuRE motifs (as a heterodimer with other NR4As), or DR5 elements (as a dimer with retinoid X receptor) (Herring et al. 2019). Through promoter binding, NR4A2 can influence cellular proliferation, inflammation, and extracellular matrix remodeling, all of which are critical to keloid biology (Herring et al. 2019; McCoy et al. 2015). However, the downstream impact of NR4A gene expression is nuanced and context-dependent, displaying both oncogenic and tumor suppressive capacity depending on the type of cancer (Crean and Murphy 2021). NR4A2 downregulation in keloid tissue with cellular hyperproliferation would indicate a tumor suppressive role in line with breast and ovarian cancers (Crean and Murphy 2021). Further, NR4A2 downregulation is associated with chronic inflammation and dysregulation, consistent with the keloid immune environment (Crean and Murphy 2021). Still, the bulk nature of RNA-Seq data precludes the identification of specific cell types linked to the NR4A2 downregulation. Further study using scRNA-Seq may help to elucidate the cell type-specific function of NR4A2 in keloids.

Additionally, IPA analysis highlighted the predicted activation of Tumor Necrosis Factor-associated genes related to tumorigenic processes in keloid tissue. While the TNF gene itself was not measured as significantly increased in KD relative to matched normal tissue, TNFR/TNF superfamily members TNFRSF9 (Log2FC = 2.30, p = 0.037) and TNFSF4 (Log2FC = 1.19, p = 0.017) were increased in keloid tissue. TNFSF4 is known to promote immune activation and fibroblast-immune crosstalk, most notably with activated CD4 T cells leading to NF-kB signaling that enhances cell survival in those T cells (Croft 2010; Sadrolashrafi et al. 2024). A prior study found that TNF-α itself was not increased in keloid tissue, but rather the soluble receptor (sTNFR1), thus increasing TNF-α sensitivity and promoting sustained/excessive proliferation of keloid fibroblasts (Li et al. 2021).

Pathway-level analyses revealed predicted activation of multiple fibrotic, immune, and tumor-related pathways. Collagen biosynthesis, degradation, and ECM organization are well documented as contributors to keloid pathogenesis (Limandjaja et al. 2020; Ogawa 2017). Further, keloids inherently develop from abnormal wound scarring, thus signaling related to the S100 family, which are damage-associated molecular patterns (DAMPs) (Gonzalez et al. 2020), eicosanoids (mediators of inflammation) (Yasukawa et al. 2020), and abnormal platelet homeostasis (Yan et al. 2021) would be expected. Moreover, integrins that mediate cell-cell surface interactions were also predicted to be activated, likely leading to increased immune cell infiltration, a hallmark of keloid tissue (Lee et al. 2023). Lastly, the HOTAIR regulatory pathway involving an oncogenic long non-coding RNA was predicted activated in keloid tissue and is known to promote tumor growth and epithelial to mesenchymal transition (Balas and Johnson 2018).

Beyond the predicted upregulated pathways related to fibrosis and immune regulation summarized above, multiple malignant cancer-associated pathways were also implicated in KD. While IPA is well known to have an over-abundance of cancer-related data, connections between KD and cancer are indeed present within the literature (Tan et al. 2019; Ud-Din and Bayat 2020). Both keloids and cancerous tumors have been described as wounds that fail to heal; however, keloids never undergo malignant transformation (Tan et al. 2019; Ud-Din and Bayat 2020). Despite this distinction, both conditions share a fibrotic tissue microenvironment, activation of epithelial to mesenchymal transition pathways, and are influenced by tissue infiltrating lymphocytes (Lee et al. 2023; Yang et al. 2018; Zhang et al. 2023a). For example, a predominance of M2 macrophages, known to be both profibrotic and pro-cancerous, is associated with poor clinical outcomes in both KD and triple negative breast cancer (Koru-Sengul et al. 2016; Nangole et al. 2021). Our findings similarly indicate increased M2 macrophages in keloid tissue relative to MANST. Additionally, recent data indicate that the propensity to develop KD is associated with increased risk of a malignant cancer diagnosis (Lu et al. 2021). Given the similarities between KD and cancer, continued study of keloid pathogenesis may offer insights into the early, pre-malignant processes that contribute to cancer development.

To address the underrepresentation of African Americans in molecular keloid research, we deliberately sampled African American patients; yet our limited sample size constrained statistical power to fully evaluate potential race-specific cell type abundances and gene expression differences. Larger multi-population studies are needed to evaluate whether racial differences could influence future KD precision medicine therapeutics. Still, ssGSEA-calculated enrichment scores enabled us to assess fibroblast subtype and immune cell contributions to gene expression in keloid tissue and MANST. This allowed us to partially disentangle the effects of fibroblast subtype and immune cell abundances from bulk tissue gene expression differences. However, our findings were limited since differential gene expression could not be resolved to specific cell type populations, a limitation readily addressed by scRNA-Seq and spatial transcriptomic/proteomic approaches. Despite this limitation, we successfully validated our primary findings in a publicly available bulk RNA-Seq dataset of African American keloid tissue with MANST from Zhu et al. (Zhu et al. 2023). While this dataset was smaller than ours (N=8 patients), it was the only comparable dataset in the public domain. Considering that this dataset was composed of keloid tissue and MANST from the torso (abdomen, chest, and back) and extremities (arm), it suggests that our findings are generalizable to keloids at other anatomical locations beyond the head and neck region.

The etiology of KD is clearly multifaceted, with both fibrotic and inflammatory contributions. Our study revealed differentially abundant cell types contributing to both components. By accounting for this cellular heterogeneity across tissue types, we uncovered gene expression differences that may more accurately reflect differential gene expression underlying KD, as opposed to genes simply marking differentially abundant cell types. Further, tissue-specific fibroblast-immune cell associations such as mesenchymal fibroblast-activated mast cell or secretory reticular fibroblast-activated NK cell may underpin KD progression since they were specific to keloid tissue. It would be a logical next step to evaluate whether variation in these keloid tissue-specific cell type enrichments, pairwise cell type correlations, and gene expression also impact KD clinical outcomes, including symptoms and keloid recurrence.

## MATERIALS & METHODS

### Study recruitment and tissue preparation

This study was conducted within Henry Ford Health (HFH), a large health care system covering southeastern and southcentral Michigan. The cohort was composed of 14 patients presenting for surgical removal of keloids located in the head and neck region between 2011 and 2017. Patient race was self-reported. Keloid diagnosis was defined clinically by the presence of a scar which had grown past the initial injury. Keloid tissue was surgically removed with a visually negative margin around the keloid with histological confirmation. Visually adjacent normal skin was also removed for comparison as matched adjacent normal skin tissue (MANST). Tissue specimens were placed in RNAlater solution then preserved in liquid nitrogen at −80 °C until sequencing. HFH Institutional Review Board approved the protocol under study number 6745. All research was conducted in accordance with the Declaration of Helsinki (2013).

### Bulk RNA extraction, sequencing, and data processing

Total RNA was extracted from tissue specimens using the Ambion RiboPure^TM^ kit. Strand-specific sequencing libraries were constructed using the TruSeq Stranded Total RNA Library Prep Kit with Ribo-Zero Globin (Illumina, San Diego, CA). RNA-Seq was performed on an Illumina NovaSeq 6000 with a target of 50 million, 150 base pair paired-end reads per sample. Raw sequencing data were input into the nf-core/rnaseq pipeline (v3.21.0) with default options except for “skip_markduplicates” set to true (Ewels et al. 2020). The Homo sapiens GRCh38.115 genome was used as reference. Briefly, reads were preprocessed to auto-infer standedness, check quality, and trim/remove adapters/contaminants as well as rRNA. Reads were then aligned with STAR (Dobin et al. 2013) and quantified with Salmon (Patro et al. 2017). Merged raw estimated gene counts for all samples were filtered to those with expression greater than 25 counts in at least 50% of the samples (14/28), leaving 15,753 genes for subsequent analyses.

### Cell type enrichment scoring

The rank-based single-sample gene set enrichment analysis (ssGSEA) algorithm was applied to the keloid and MANST RNA-Seq filtered raw estimated count data to calculate the enrichment score for each fibroblast subtype in each of the keloid tissue and MANST specimens. The five fibroblast subtypes (and the corresponding differentially expressed genes) defined by Sole-Boldo et al. (Solé-Boldo et al. 2020) were used as the reference for the ssGSEA scoring. Briefly, we first filtered DEGs in the Sole-Boldo ‘Supplementary Data 3’ table by significance (adjusted p-value < 0.05). Next, gene sets were defined as the top 50 DEGs for each fibroblast subtype sorted by highest average log fold change (**Table S7**). Finally, ssGSEA scores based on the gene sets were converted to Z-scores to allow for comparison of subtype enrichment across samples. The same ssGSEA enrichment scoring was done for immune cells, with gene sets for each immune cell type derived by assigning genes to cell type based on their maximum value in the LM22 signature matrix file available from CibersortX (Newman et al. 2019) (**Table S7**).

### Differential fibroblast subtype and gene expression testing between keloid and MANST

Differential expression testing of individual 1) fibroblast subtype enrichment scores and 2) genes between keloid and MANST was performed using mixed-effects models to account for the paired study design, with keloid and MANST samples obtained from the same patient. Fibroblast subtype enrichment scores were evaluated with linear mixed-effects models, whereas DEGs were evaluated using negative binomial generalized linear mixed-effects models with a log link. In both cases, tissue status (i.e. keloid/MANST) was treated as a fixed effect, and patient identity was included as a random effect. Further, for DEG analyses, raw counts for each gene were modeled as the outcome, and an additional edgeR-based (v4.6.3) offset term was used to account for differential sample library size (Robinson et al. 2010). Nominal p-values for DEGs were adjusted for multiple testing using the Benjamini and Hochberg false discovery rate (FDR) approach (Benjamini and Hochberg 1995).

### Ingenuity Pathway Analysis of DEGs

To interpret biological pathway enrichment of DEGs (keloid/MANST and between pairs of keloid subgroups), QIAGEN Ingenuity Pathway Analysis (IPA) (QIAGEN Inc., https://digitalinsights.qiagen.com/IPA) was employed; accessed May 2026 (Krämer et al. 2014).

DEGs were identified by Ensembl ID and HUGO gene symbol for an analysis of IPA canonical pathways, which are well-characterized metabolic and cell signaling pathways. Significant pathway enrichment was determined by a Fisher’s Exact Test, with FDR adjustment of resulting p-values to account for multiple testing. Further, Z-scores predicting the overall activation/inhibition state of each pathway were determined based on how closely expression patterns match the expected pattern for a given pathway (Krämer et al. 2014).

### Data visualization

The forest plot was generated with the R package forestplot (V3.1.7) (Gordon and Lumley 2026). The dotchart, histogram with density curves, and barplot were generated with the R package ggplot2 (V4.0.1) (Wickham 2016). All analyses were performed in R V4.5.0 (R Core Team 2025).

### Validation

To externally validate our findings in an independent keloid dataset, bulk RNA sequencing data from Zhu et al. (Zhu et al. 2023) were downloaded from the NCBI sequence read archive (Bioproject PRJNA905476). Consistent with our patient-matched tissue study design, this dataset included RNA-Seq data from paired untreated keloid tissue and MANST (non-lesional) specimens from 8 African American patients. The corresponding fastq RNA-Seq data files were re-processed with the nf-core/rnaseq pipeline (as described above) to generate a gene level count table. Using these data, cell type enrichment scoring and tissue type association analyses (differential cell type enrichment, cell-cell associations, and gene expression) were all performed using the same methods as described above.

## ETHICS

This study was performed in accordance with the Declaration of Helsinki. Collection of human tissue samples for this study was approved as part of the study protocol. This human study was approved by the Henry Ford Health Institutional Review Board - approval: 6745. All adult participants provided written informed consent to participate in this study.

## DATA AVAILABILITY

The RNA-sequencing data generated in this study have been deposited in the Gene Expression Omnibus (GEO) under accession number GSE344399. All other relevant data supporting the findings of this study are available from the corresponding author upon reasonable request.

## ACKNOWLEDGEMENTS

We would like to thank the patients who participated in the study. This work was supported by funding from Henry Ford Health Game on Cancer (H10338), National Institutes of Health (K08GM128156), and American Academy of Facial Plastic and Reconstructive Surgery (350651/F19538). Additionally, IA received funding from NIH/NIAMS (R01AR083553).

## AUTHOR CONTRIBUTIONS

Conceptualization: JJP, AML, LRJ

Data Curation: JJP, MP, AML, LRJ

Formal Analysis: JJP, MP, AML, IML

Funding Acquisition: AML, LRJ

Investigation: JJP, MP, AML, LRJ

Methodology: JJP, AML

Project Administration: MN

Resources: LRJ

Software: JJP, MP, IML

Supervision: IA, AML, LRJ

Validation: LRJ

Visualization: JJP

Writing – Original Draft Preparation: JJP, MP, AML, LRJ, IA

Writing – Review & editing: JJP, MP, MN, IML, IA, LH, DC, RF, DBVC, CDGS, AML, LRJ

## DECLARATION OF GENERATIVE AI USE

The author(s) did not use AI/LLM in any part of the research process and/or manuscript preparation.

## SUPPLEMENTARY MATERIAL

**Figure S1.**
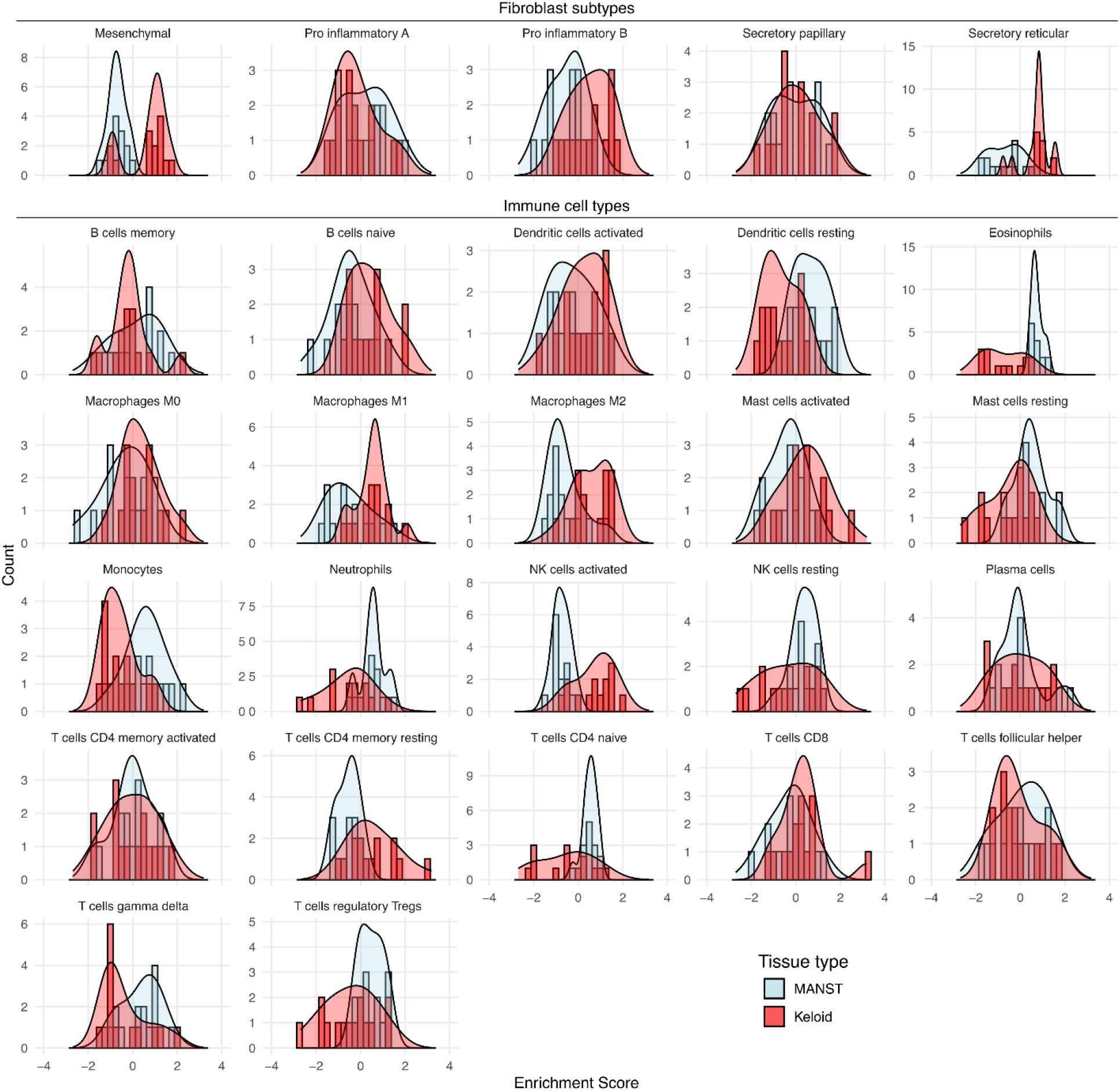
Enrichment score distributions for fibroblast subtypes and immune cells across tissue type. Histograms plotted with density curves colored by tissue type to highlight enrichment score distributions across tissue type.

**Figure S2.**
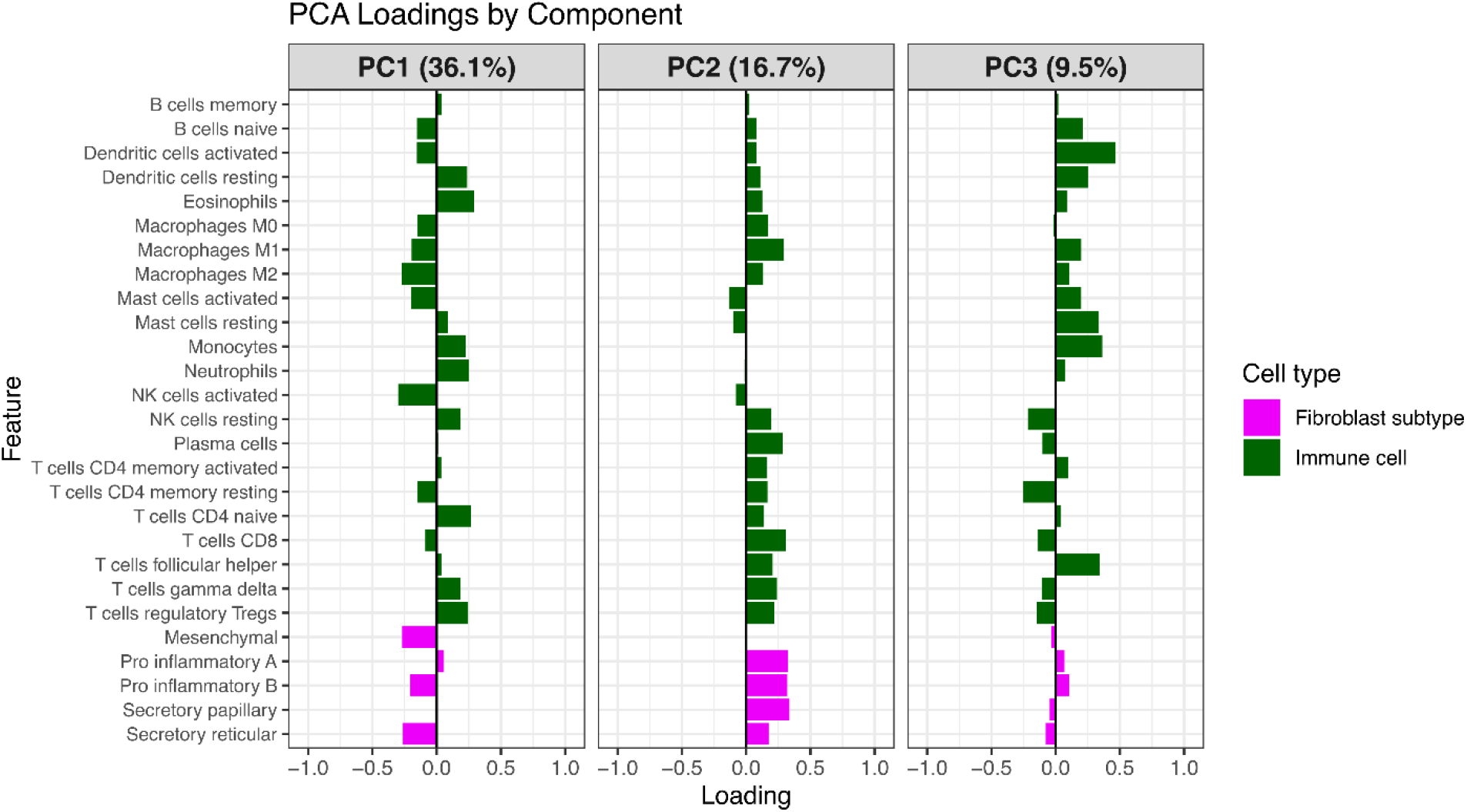
Fibroblast subtype and immune cell enrichment score loadings by principal component. Loadings for the first 3 principal components of a PCA based on all 27 enrichment scores. Loadings were grouped by cell type and sorted alphabetically within each grouping.

### Supplementary Table Legends

**Table S1. Validation of tissue type enrichment score associations from the HFH cohort in the Zhu et al. dataset.** Both fibroblast (n=3) and immune cell (n=14) enrichment score associations meeting the significance threshold (p<0.05) in the HFH dataset were evaluated for tissue type association in the Zhu et al. dataset. The resulting model coefficients, 95% confidence intervals, and p-values were juxtaposed with the corresponding HFH associations.

**Table S2. Association between fibroblast subtype and immune cell type enrichment scores stratified by tissue type with interaction.** Associations between enrichment scores for fibroblast subtypes and immune cell types were evaluated with a tissue type-stratified linear mixed effects model accounting for patient identity as a random effect along with a tissue type interaction term. Significance of the interaction term between tissue type and fibroblast subtype is included to the right of the tissue type-specific results. Additionally, differential correlation patterns were indicated for each fibroblast subtype-immune cell association defined as β_Keloid_ > β_MANST_ (Pattern 1) and β_Keloid_ < β_MANST_ (Pattern 2).

**Table S3. Validation of tissue type-specific fibroblast subtype-immune cell associations from the HFH cohort in the Zhu et al. dataset.** Tissue type-specific fibroblast subtype-immune cell associations meeting the significance threshold (interaction p-value < 0.1) in the HFH dataset were evaluated in the Zhu et al. dataset. The resulting model coefficients, 95% confidence intervals, and p-values were juxtaposed with the corresponding HFH tissue type-specific cell-cell associations. Consistency in the coefficient patterns (Pattern 1 - Coef_Keloid_ > Coef_MANST_; pattern 2 - Coef_Keloid_ < Coef_MANST_) across datasets was also evaluated.

**Table S4. Gene expression associations with tissue type.** Differential gene expression analysis results between keloid tissue and MANST. Negative binomial mixed-effects model was adjusted for the first 3 PC’s of the cell type enrichment score PCA. P-values were adjusted for false discovery rate.

**Table S5. Validation of tissue type gene associations from the HFH cohort in Zhu et al. dataset.** Gene associations with tissue type were filtered to those with an FDR-adjusted p-value < 0.05 in the HFH dataset. Gene expression associations with tissue type were evaluated in the Zhu et al. dataset. Directionality of the log_2_ fold-changes were compared across datasets.

**Table S6. IPA pathway enrichment by differentially expressed genes across tissue type.** Pathway gene enrichment significance was determined by p value of differentially expressed gene overlap with total genes in IPA pathways. P values were adjusted by false discovery rate correction. Proportion column represents the percentage of DEGs corresponding to the pathway divided by the total number of genes in the IPA pathway.

**Table S7. Gene sets for fibroblast subtypes and immune cell types.** Fibroblast subtype marker gene lists from Sole-Boldo et al. were filtered to genes in the bulk RNA-seq dataset. Mitochondrial-related genes were also filtered out. The top 50 by average logFC with adjusted p-values < 0.05 were selected per fibroblast subtype. Immune cell type gene sets were determined by assigning genes to cell types based on their highest expression value in the LM22 signature matrix.

